# Mixed Origins: HIV gp120-specific memory develops from pre-existing memory and naïve B cells following vaccination in humans

**DOI:** 10.1101/2021.09.01.458551

**Authors:** Madhubanti Basu, Michael S. Piepenbrink, Christopher Fucile, Catherine A. Bunce, Li-Xing Man, Jane Liesveld, Alexander F. Rosenberg, Michael C. Keefer, James J. Kobie

## Abstract

The most potent and broad HIV envelope (Env)-specific antibodies often when reverted to their inferred germline versions representing the naïve B cell receptor, fail to bind Env suggesting that the initial responding B cell population is not exclusively comprised of a naïve population, but also a pre-existing cross-reactive antigen-experienced B cell pool that expands following Env exposure. Previously we isolated gp120-reactive monoclonal antibodies (mAbs) from participants in HVTN 105, an HIV vaccine trial. Using deep sequencing VH-lineage tracking we identified several of these mAb lineages in pre-immune peripheral blood. Several of these pre-immune lineages also persisted in the bone marrow, including CD138+ long-lived plasma cell compartment, ∼7 months after the final vaccination. The majority of the pre-immune lineage members included IgM, however IgG and IgA members were also prevalent and exhibited somatic hypermutation. These results suggest that vaccine-induced gp120-specific antibody lineages originate from both naïve and cross-reactive memory B cells.

## Introduction

Despite increased access to antiretrovirals for treatment and prevention, HIV-1 is still a major health burden with ∼1.5 million new infections and ∼680,000 HIV-related deaths globally in 2020 (WHO, 2020). Thus, development of a safe and effective preventive HIV vaccine remains a global priority. HIV-1 Envelope (Env) glycoprotein is the primary target of the humoral response to HIV-1 infection and the antibody (Ab) response has been correlated with vaccine mediated protection from HIV infection (Chung et al., 2015; Haynes et al., 2012; Karasavvas et al., 2012; Yates et al., 2014; Zolla-Pazner et al., 2014a; Zolla-Pazner et al., 2013). Although Env is highly immunogenic and readily induces Abs, Ab-mediated protection from HIV appears to be conferred if at all, by only a minor subset of Ab clones from the overall Env-specific immunoglobulin (Ig) repertoire. Approximately 10-50% of people living with HIV (PLWH) within several years of infection, develop Abs with tremendous breadth able to neutralize diverse isolates, although typically not their own contemporary isolates (Bonsignori et al., 2017; Hraber et al., 2014), indicating that such broadly neutralizing antibodies are capable of being induced, and that if recapitulated by vaccination would likely confer protection from infection.

Numerous HIV-1 broadly neutralizing monoclonal antibodies (bnmAbs) have been isolated from PLWH and are the most well characterized of HIV-1 specific mAbs including extensive deep-sequencing analysis of their clonal lineages. These bnmAbs typically have extensive somatic hypermutation and common molecular features in the heavy and light chain variable regions. Some of these bnmAbs share very specific structural features like gene usage and complementarity-determining region 3 (CDR3) lengths, particularly observed among those targeting the CD4 binding site, referred to as VRC01-class bnmAbs (West et al., 2012; Wu et al., 2015). Many of these bnmAbs when reverted to their presumed germline or naïve sequence fail to bind HIV Env (Ahlers, 2014; Hoot et al., 2013; Xiao et al., 2009b) raising the possibility that they may arise from naïve B cells that responded initially to a different antigen, acquiring cross-reactivity with HIV Env sufficient to respond HIV infection, and then subsequently developed increasing affinity to HIV Env as part of their extensive somatic hypermutation process in response to chronic viremia. Recently, the development of ‘germline-targeting’ immunogens has showed their potential for binding predicted germline versions of bnmAbs, activating transgenic B cells and generating antibody responses in B cell receptor (BCR) transgenic mouse models (Abbott et al., 2018; Briney et al., 2016; Escolano et al., 2016; Havenar-Daughton et al., 2018; Huang et al., 2020; Jardine et al., 2013; Jardine et al., 2015; Lin et al., 2020). Havenar-Daughton et al, 2018 have used the eOD-GT8 immunogen successfully to isolate genuine human naïve B cells *ex vivo*, showing some promise for the ‘germline-targeting’ vaccine development concept (Havenar-Daughton *et al*., 2018). Subsequently, the eOD-GT8 60mer immunogen is in the early stages of clinical testing to determine the potential of germline targeting approaches for stimulating the human B cell repertoire (ClinicalTrials.gov, 2021).

Several HIV-1 vaccine efficacy trials have been completed so far although no vaccine has been licensed to date (Ng’uni et al., 2020). RV144 is the only immunogen-based preventive HIV vaccine trial that has demonstrated moderate protection so far, however this regimen did not induce a substantial neutralizing antibody response (Rerks-Ngarm et al., 2009). More recently two phase 2b Antibody Mediated Prevention (AMP) trials with the VRC01 bnmAb infusion in high-risk populations have found that the VRC01 was able to block acquisition of only ∼30% of the circulating strains, however, it was 75% effective at preventing acquisition of VRC01-sensitive strains (in vitro IC_80_ <1 µg/ml) (Corey et al., 2021; Edupuganti et al., 2021; Mgodi et al., 2021). Although VRC01 could not prevent overall HIV-1 acquisition more effectively than placebo, these ‘proof-of-concept’ studies indicated the potential of bnmAbs to prevent HIV infection. All these studies highlight the need to understand the mechanisms of bnAb formation in order to develop an effective HIV-1 vaccine (Edupuganti *et al*., 2021; Mgodi *et al*., 2021).

Several findings from PLWH support that one of the possible developmental pathways for HIV-1 Env antibodies involve poly-reactive B cells that cross-react with self and/or microbial antigens, particularly those that target the gp41 component of Env (Mouquet and Nussenzweig, 2012; Planchais et al., 2019; Trama et al., 2014). Additionally, Williams et al. reported that following HIV vaccination, a gp41-reactive mAb lineage which cross-reacts to gut residing *E. coli* RNA polymerase developed. They further suggested that such cross-reactive populations may result in an immunodominance that minimizes the development of other Env-specific B cell responses to vaccination (Williams et al., 2015). In general, HIV-1 Env vaccine containing both gp120 and gp41 primarily induces an intestinal microbiota derived cross-reactive gp41-dominant response along with a low frequency of gp120 response (Williams *et al*., 2015) whereas immunization with gp120-only results in a comparatively higher frequency of gp120-specific memory B cell response than the response that is induced from gp140 or gp160 vaccination (Moody et al., 2012; Wiehe et al., 2014). Moreover, several lines of evidence indicate that the precursors of Env-specific mAb lineages can be identified in the naïve B cell repertoire (Havenar-Daughton *et al*., 2018) and some antibodies with bnAb activity can also arise from naïve B cells (Krebs et al., 2019).

Previously we isolated 66 gp120 reactive mAbs (Basu et al., 2020) from HVTN 105 trial participants (Rouphael et al., 2019), a phase I trial testing AIDSVAX B/E (gp120 protein) that was used in RV144, combined with a DNA immunogen. In order to determine the origin of these mAb producing B cells, we used deep sequencing-based VH lineage tracking and identified several of these mAb lineages in peripheral blood prior to vaccination (pre-immune). The majority of the pre-immune gp120-reactive lineages included IgM, however IgG and IgA members were also predominant prior to vaccination, Additionally the majority of the pre-immune lineage members exhibited somatic hypermutation prior to vaccination, suggesting that the vaccine-induced gp120-specific antibody lineages originated from both naïve B cells and cross-reactive memory B cells.

## Results

### Identification of HIV-1 Env-specific B cells in pre-immune blood

We have previously observed the presence of HIV-1 Env gp140 reactive B cells in healthy participants both within the CD27-naïve B cell and CD27+ memory B cell compartments (**Figure 1A**), suggesting the presence of pre-existing non-naïve HIV-1-Env reactive B cell populations. To characterize this pre-immune HIV-1 Env reactive B cell compartment further we collected peripheral blood samples from 21 healthy participants of the HVTN 105 HIV-1 vaccine clinical trial before and after immunization (**Figure 1B**). The participants received 4 vaccinations over a period of 6 months with different combinations of AIDSVAX B/E bivalent gp120 protein, which includes gp120 MN.B and gp120 A244.AE proteins, and DNA-HIV-PT123, consisting of plasmids expressing 96ZM651.C gp140, ZM96.C gag, and CN54.C pol-nef (Rouphael *et al*., 2019). Human gut microbiota reactive antibodies are known to recognize gp41 subunit of the HIV-1 Env (Cram et al., 2019; Planchais *et al*., 2019; Rouphael *et al*., 2019; Williams et al., 2018; Zolla-Pazner et al., 2014b). Therefore, we decided to isolate pre-immune HIV-1-Env-specific (gp120pos) B cells using gp120 MN.B protein that was a part of HVTN 105 vaccine. The remaining fraction of B cells (gp120neg) was used for comparison.

**Figure 1.**
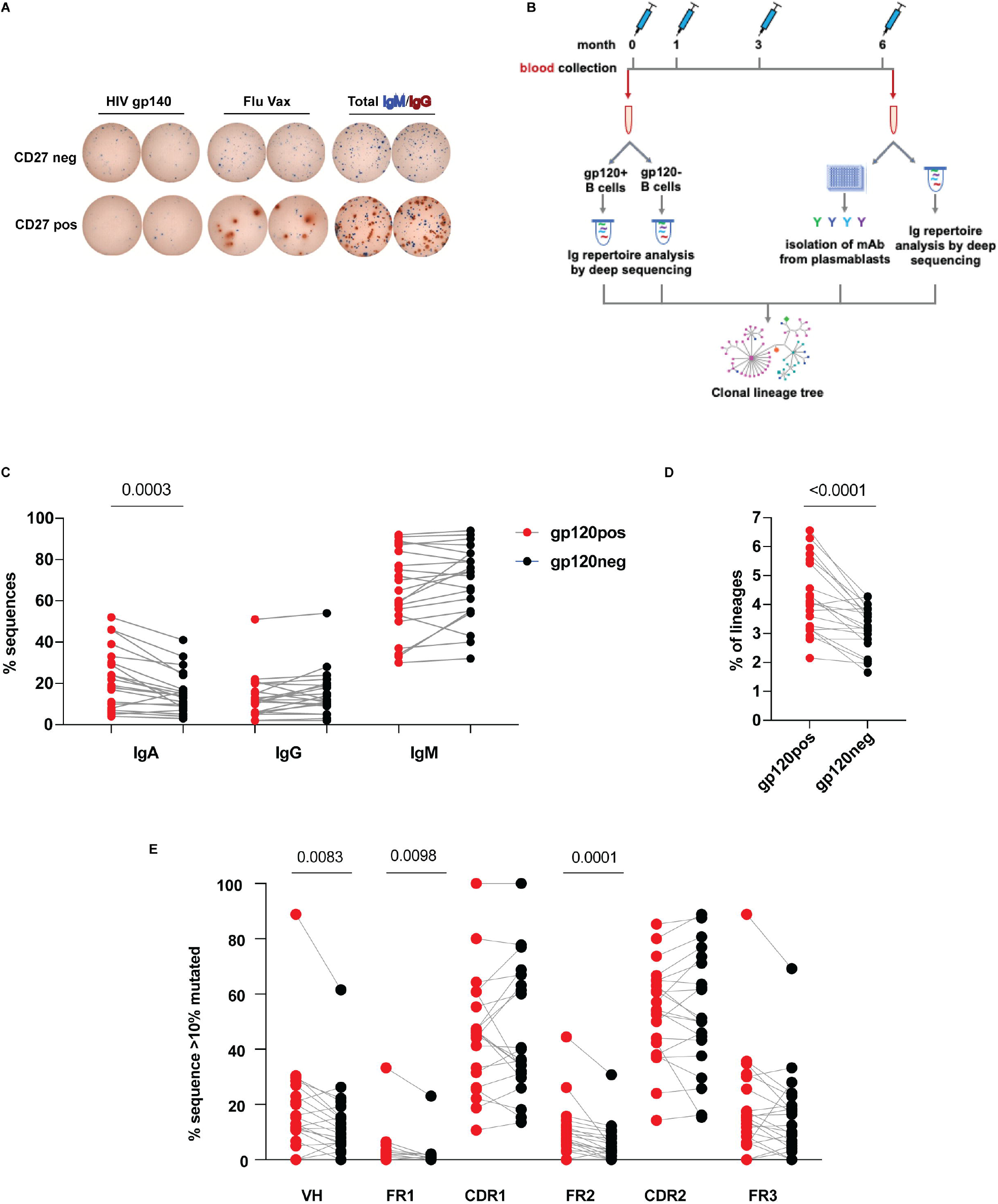
Characterization of pre-immune B cell repertoire in blood. (A) Representative ELISpot showing pre-immune CD27pos (memory) and CD27neg Env-reactive B cells in HIV negative healthy subjects (Blue=IgM, Red=IgG). (B) Timeline showing the collection of peripheral blood at baseline (Month 0) and seven days after final immunization (M6) from HVTN 105 participants (n = 21) who received AIDSVAX B/E Protein and DNA-HIV-PT123 plasmid immunizations intramuscularly at months 0, 1, 3, and 6. (C-F) Frequency of (C) IgA, IgG and IgM, (D) lineages with long HCDR3 (≥22 amino acid), (E) expanded lineages with ≥ 10% average mutation from germline in gp120pos and gp120neg B cell fractions.

### Characterization of pre-immune B cell repertoire in blood

To understand the molecular features of HIV-1 Env specific pre-immune BCR repertoire we analyzed characteristics of the heavy chain variable region by deep sequencing of these gp120pos and gp120neg B cells from all 21 participants. Comparing the isotype distribution (**Figure 1C**), we found IgM was the predominant isotype in both the gp120pos and gp120neg repertoires overall, as expected since naïve IgM+ B cells comprise the majority of the peripheral B cell compartment in healthy individuals. Non-IgM isotypes (i.e. IgG or IgA) were also substantially present in gp120pos (ranging from 3.8-52% for IgA and 1.5-50.9% for IgG) as well as in gp120neg (ranging from 2.7-41.5% for IgA and 1.9-53.7% for IgG) in pre-immune blood. However, on the individual participant basis, higher frequencies of IgM (in 15/21; 71% of the participants) and IgG (in 12/21, 57%) were found in gp120neg compared to gp120pos. In contrast, the gp120pos fraction was significantly enriched (p=0.0003) for IgA with a higher IgA proportion in 86% of the participants (18/21) compared to their gp120neg population.

The CDR3 is the most variable region of the antibody molecule and a major determinant for antigen recognition and binding. Among various known characteristics, the presence of long HCDR3, as defined as > 22 aa, is a common feature for some well-characterized bnmAbs against HIV-1, and several Env-reactive mAbs isolated from HIV vaccine trials also possess this feature (Basu *et al*., 2020; Easterhoff et al., 2017). We searched for this trait in the pre-immune peripheral blood B cells and found significant (p<0.0001) enrichment of lineages with long HCDR3 in the gp120pos over gp120neg fraction in 17 of the 21 (∼81%) participants (**Figure 1D**).

Next, we wanted to assess the presence of somatically mutated Abs in pre-immune repertoire as an indication of antigen experience. Since the predominant population in peripheral blood of healthy is expected to be naïve B cells which lack somatic hypermutation, BCR sequences with > 10% nt mutations compared to germline were analyzed in order to focus on the non-naive population only. There was no significant difference in overall VH mutation level between the gp120pos and gp120neg non-naïve B cell fractions (not shown). Assessment of expanded clonal lineages, those that predominate the repertoires, revealed that there was significantly (p=0.0083) higher frequency of mutated (>10% nt) VH sequences in gp120pos (**Figure 1E**). This increased VH mutation in gp120pos compared to gp120neg was most evident in framework region 1 (FR1) (p=0.0098) and FR2 (p=0.0001). Together these results indicate the pre-immune gp120pos compartment includes IgM, IgG, and IgA B cells with proportionally more IgA, clones with longer HCDR3, and increased somatic hypermutation in FR1 and FR2.

### Expansion of HIV-1 Env specific pre-immune B cells following HIV Env vaccination

The pre-immune HIV Env-reactive B cell fraction is assumed to not have had any prior exposure to HIV Env and is anticipated to have a certain degree of representation in the resulting HIV Env vaccine-induced B cell repertoire. Previously, for the HIV Env vaccine phase 1 trial HVTN 105, BCR repertoires from day 7 post-vaccination peripheral blood (D7), expected to be dominated by vaccine-induced IgG plasmablasts, were sequenced (Basu *et al*., 2020). We sought to understand the degree of overlap between the gp120pos pre-vaccination and the vaccine-induced BCR repertoire. Nine out of 11 participants for which D7 samples were analyzed had post-vaccination lineages that could be traced back to the HIV-1 Env specific pre-immune B cell compartment. However, as expected, due to the large size and diversity of the repertoire and limitation of sequencing depth, only a few lineages were shared between the pre- and post-vaccination samples. To understand whether these shared pre-immune lineages originated from the naïve compartment or not, pre-immune sequences having <5% nucleotide mutations from germline in the variable gene were considered derived from naïve B cells, and those with >10% nucleotide mutations from germline in the variable gene were considered derived from non-naïve B cells. Although a higher number of shared lineages between D7 and the pre-immune gp120pos were found in the naïve than the non-naïve compartment (**Figure 2A**), overall, they represented a small frequency (< 0.5%) of the naïve population of gp120pos pre-immune. In contrast, the shared lineages between D7 and the gp120pos pre-immune were found in a proportionally significantly (p=0.0244) higher frequency (highest ∼2.5%) of the non-naïve compartment of the gp120pos pre-immune (**Figure 2B**).

**Figure 2.**
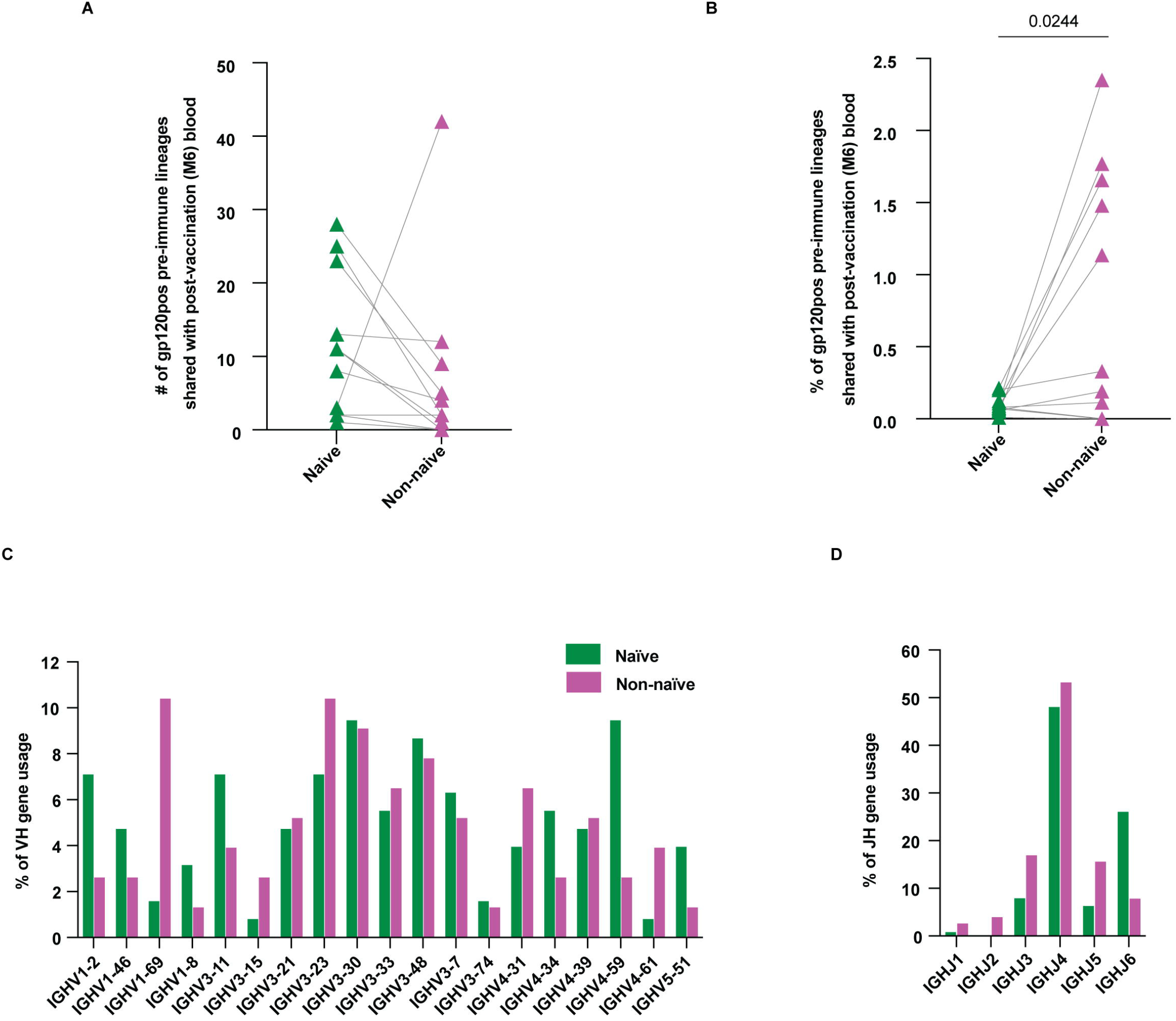
Expansion of HIV-1 Env specific pre-immune B cells following HIV Env vaccination. gp120pos pre-immune B cells that shared lineages with M6 peripheral blood that was obtained D7 after final immunization, were split into naïve (<5% nucleotide mutations from germline in the variable gene) and non-naïve (>10% nucleotide mutations from germline in the variable gene) compartments and characterized. (A) Number of lineages, (B) percentage of lineages, (C) VH and (D) JH gene usage.

Among the lineages that are shared between pre-immune and post vaccination (D7), there was a higher frequency of VH1-2 and VH4-59 lineages in the naïve than in non-naïve pre-immune lineage members (**Figure 2C**). Contrastingly, the frequency of VH1-69 lineages was higher in non-naïve (10.4%) than in naïve (1.57%) pre-immune members. Among the JH gene families, we didn’t find any difference except lower JH6 usage in naïve than in non-naïve population (**Figure 2D**). Together these results suggest that the non-naïve gp120pos pre-immune compartment is preferentially responsive to this HIV vaccine than the naive gp120pos pre-immune compartment. These results further suggest VH1-2 and VH4-59 gp120pos lineages may preferentially arise from the naïve pre-immune compartment, whereas VH1-69 gp120 lineages may preferentially arise from the non-naïve pre-immune compartment following HIV Env vaccine.

### Mixed origins of vaccine-induced gp120 specific pre-immune repertoire

Observing shared lineage members in the pre and post vaccination samples, we next sought to definitively characterize the dynamics of the gp120 pre-immune lineages that respond to vaccination. Previously we isolated 66 gp120 specific mAbs from HVTN 105 participants after their last vaccination (Basu *et al*., 2020). We sought to determine whether the precursor B cells for these mAbs were present in the pre-immune B cell compartment and subsequently participated in the Env-reactive B cell response to the vaccination. We parsed our mAbs against the pre-immune deep-sequencing database using the criteria of identical VH and JH regions, HCDR3 length and >80% HCDR3 nucleotide homology, consistent with previous deep-sequencing clonal lineage analyses (Davis et al., 2020; Setliff et al., 2018; Soto et al., 2019) to identify the pre-immune clonal lineage members, and subsequently identified six lineages from three of the study participants (**Table 1**). No distinct binding or molecular features such as V or J family gene usage distinguished these mAbs from the other mAbs that were not found in the pre-immune repertoire (not shown). Five out of these six pre-immune mAb lineages were isolated from T2 and T4 group participants, which we previously reported to have vaccine-induced dominant VH1 gene usage and unsurprisingly, 4/6 lineages were also from VH1 gene family. Pre-immune members of 3/6 lineages (1114G7, 1131D12, 1095A8) were found to be IgM and IgA and/or IgG, with a minimum of 3% VH amino acid mutation from germline, indicating they developed from a non-naïve pre-immune origin. The 1098B8 lineage members were only found as IgG in pre-immune, also indicating development from a non-naïve pre-immune origin. The 1131D12 lineage members were found as IgA and somatically mutated IgM in pre-immune indicating development from a non-naïve pre-immune origin. Members of the 1131D12 lineage were also found in tonsil 7 months after final vaccination, all as IgA, indicating mucosal presence of this lineage. Among these four non-naïve pre-immune origin lineages, substantial VH somatic hypermutation (3%-35%) in the pre-immune lineage members was evident. Two of the lineages (1098B11 and 1131A5) were found as only IgM in pre-immune, including lineage members in their germline configuration, lacking any VH amino acid mutation, suggesting the 1098B11 and 1131A5 lineages developed from a naïve pre-immune origin.

**Table 1:**
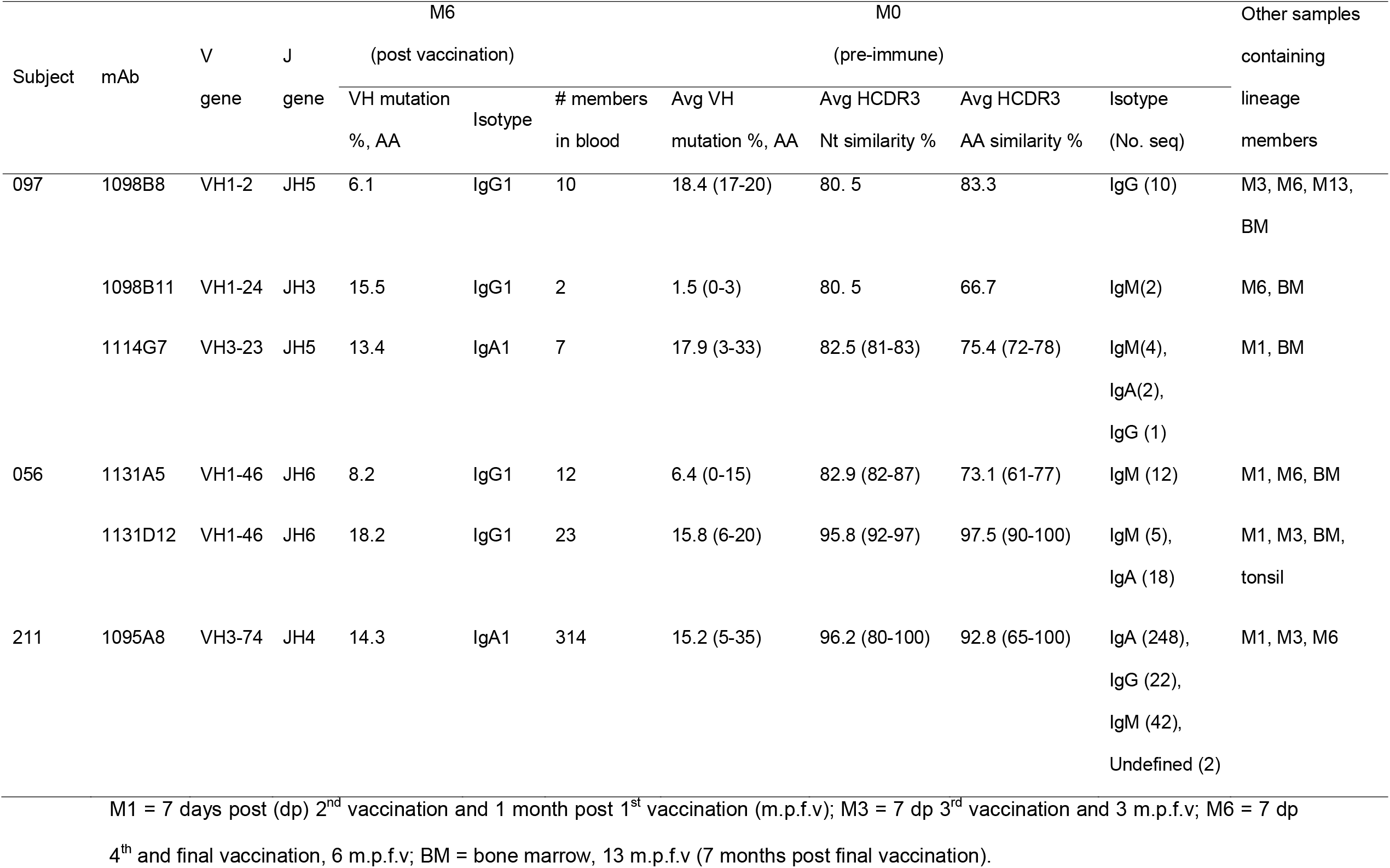
Summary of vaccine-induced gp120-reactive mAb lineages identified in pre-immune peripheral blood.

We selected three representative lineages to investigate further and constructed lineage trees from the clonally related sequences. The 1098B11 lineage (**Figure 3**), which we have previously determined targets V3 of gp120 (Basu *et al*., 2020), had the lowest somatic mutation (0-3%) observed in pre-immune members (M0) which consisted of only IgM and was considerably mutated when isolated from post-vaccination blood as an IgG mAb. This lineage is likely indicative of a naïve B cell responding to HIV gp120 proceeding through affinity maturation and class switching. Lineage members with additional somatic mutations were also found in the bone marrow 7 months after final vaccination (M13), likely representing long-lived plasma cells. The 1098B8 pre-immune lineage members (**Figure 4**) were identified as IgG and had substantial mutation from germline (17-20%). Lineage members with additional somatic mutations were also present as CD138+ long-lived plasma cells in the bone marrow 7 months after final vaccination and also in peripheral blood memory B cells. This suggests the lineage represents a non-naïve B cell responding to HIV gp120 and resulting in additional clonal expansion and somatic hypermutation. The 1095A8 lineage (**Figure 5**) includes pre-immune members that are highly somatically mutated (5-35%) and IgA, IgG, and IgM. The lineage spans multiple time points following the vaccination course, with IgA dominance in pre-immune and throughout.

**Figure 3.**
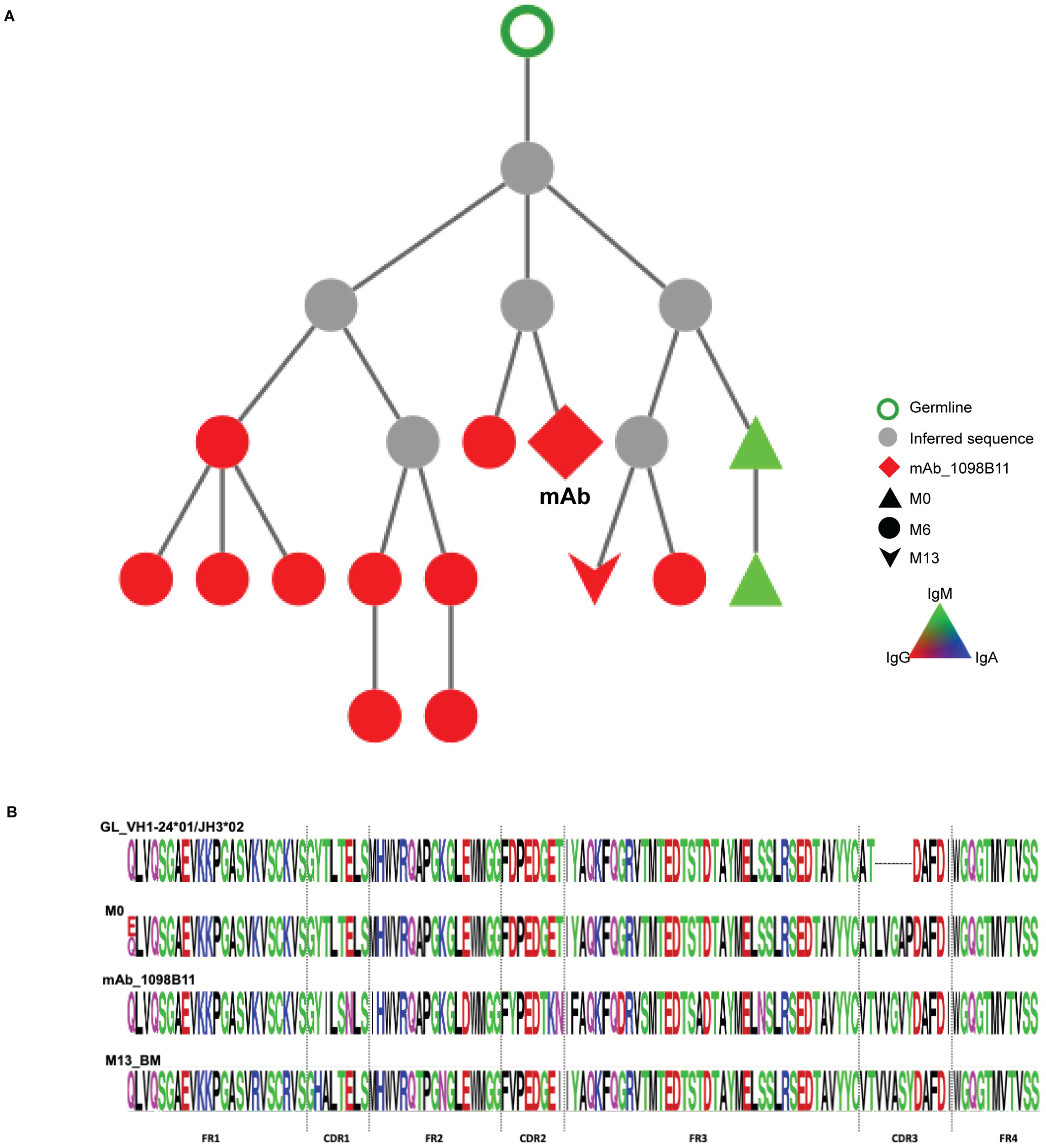
HIV-1 Env reactive 1098B11 lineage in pre-immune blood. (A) Phylogenic analysis and alignments of 1098B11 lineage. Lineage members were defined as same heavy-chain V and J gene usage, HCDR3 length, and >80% HCDR3 similarity to the mAb sequence. Individual nodes indicate identical amino acid sequences (n=1-28). (B) Alignments depict germline, pre-immune lineage members, M6 mAb, and M13 BM derived sequences.

**Figure 4.**
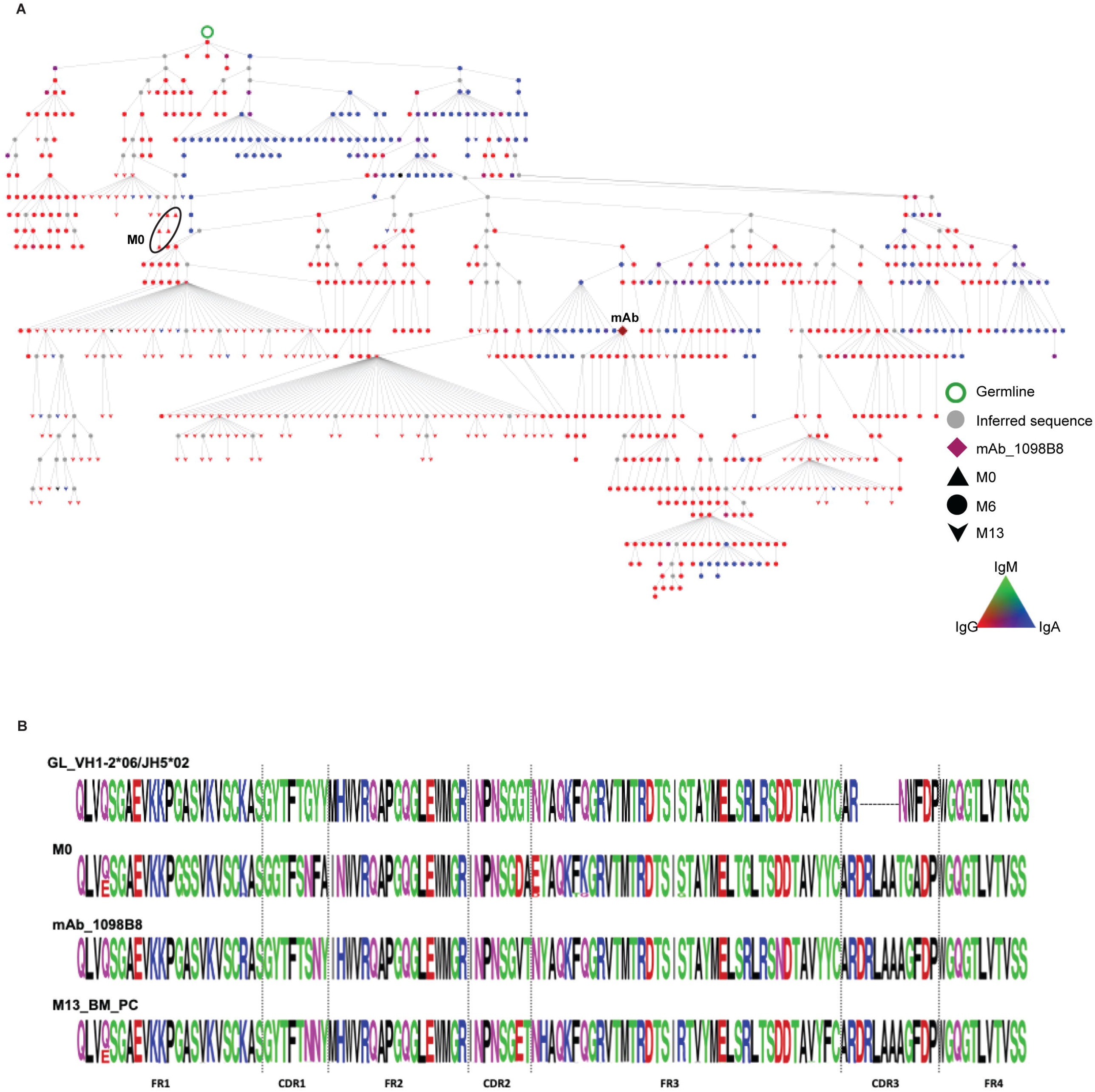
HIV-1 Env reactive 1098B8 lineage in pre-immune blood. (A) Phylogenic analysis and alignments of 1098B8 lineage. Lineage members were defined as same heavy-chain V and J gene usage, HCDR3 length, and >80% HCDR3 similarity to the mAb sequence. M13 sequences were obtained from peripheral blood, total BM and CD138+ BM PC. Individual nodes indicate identical amino acid sequences (n=1-364). (B) Alignments depict germline, pre-immune lineage members, M6 mAb, and M13 BM PC derived sequences.

**Figure 5.**
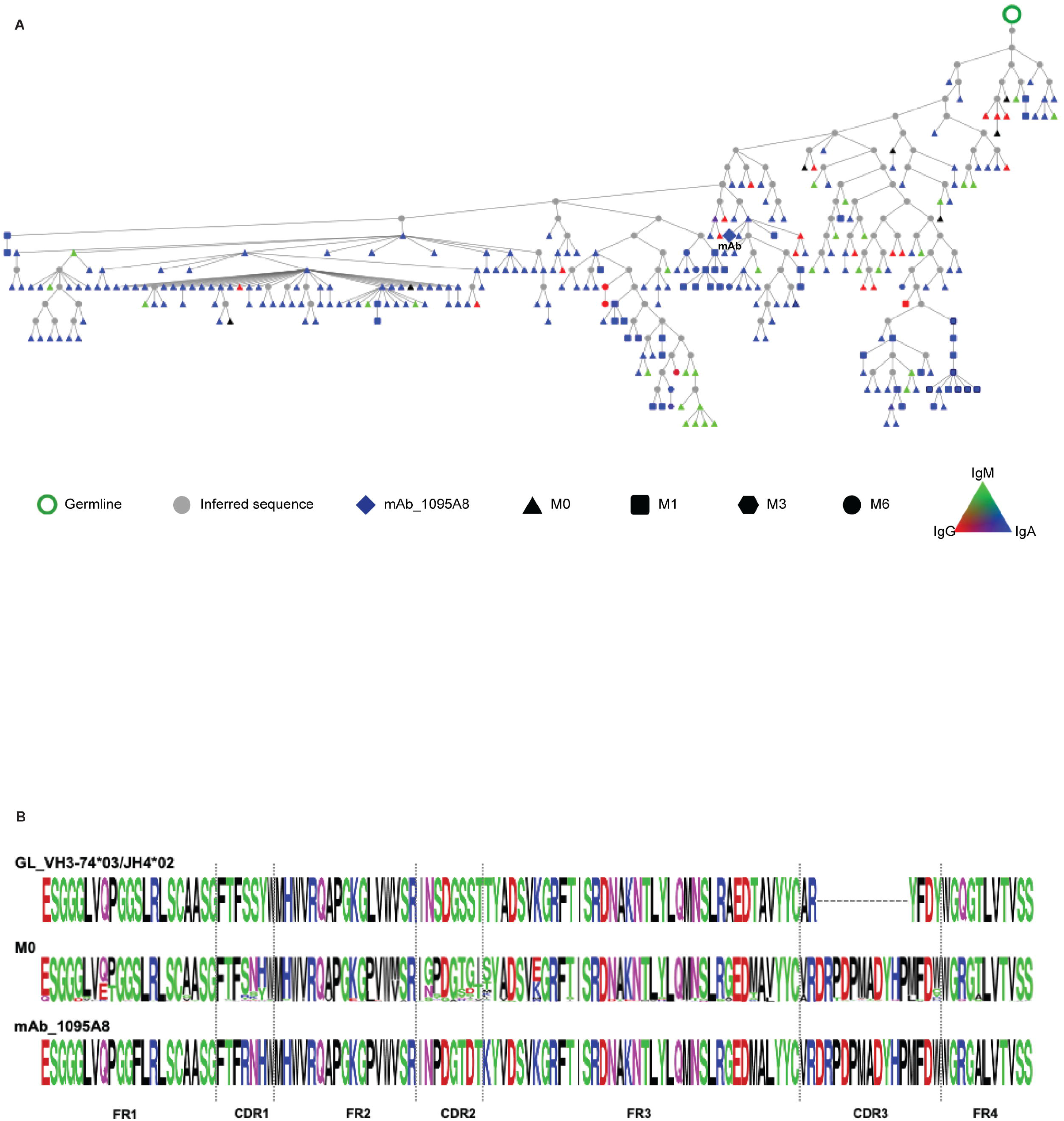
HIV-1 Env reactive 1095A8 lineage in pre-immune blood. (A) Phylogenic analysis and alignments of 1095A8 lineage. Lineage members were defined as same heavy-chain V and J gene usage, HCDR3 length, and >80% HCDR3 similarity to the mAb sequence. Individual nodes indicate identical amino acid sequences (n=2-87). Nodes with sequences from multiple time points are in black borders. (B) Alignments depict germline, pre-immune lineage members and M6 mAb sequences.

### gp120 binding by pre-immune antibodies

We tested the gp120 binding ability of the pre-immune versions of these lineages (**Figure 6**). The representatives of each pre-immune lineage and their germline versions were synthesized and complemented with the mature or the germline light chain versions. The pre-immune mAbs showed lower reactivity than the post-vaccination mature Abs against HIV-1 Env of the vaccine component strains MN.B [AIDSVAX B/E] (**Figure 6A**) and A244.AE [AIDSVAX B/E] (**Figure 6B**) but higher than their predicted germline versions. We further tested one of these lineages for the binding affinity against HIV-1 Env MN.B gp120 by surface plasmon resonance (SPR). Mature 1098B8 mAb bound gp120 with a dissociation constant (K_D_) of 236 pM demonstrating high affinity (**Figure 6C**). As expected, the pre-immune mAb showed lower but appreciable binding (K_D_ = 34.4 nM). The germline mAb showed a general lack of binding to gp120 and therefore the kinetic constant could not be similarly determined. Together these results suggest that 1098B8 developed from a non-naïve pre-immune origin as a result of an initial naïve B cell which had minimal gp120 reactivity, responding to a non-HIV Env antigen and undergoing somatic hypermutation and acquiring HIV Env cross-reactivity, and subsequently responding to the HIV Env vaccine and undergoing HIV Env-directed affinity maturation.

**Figure 6.**
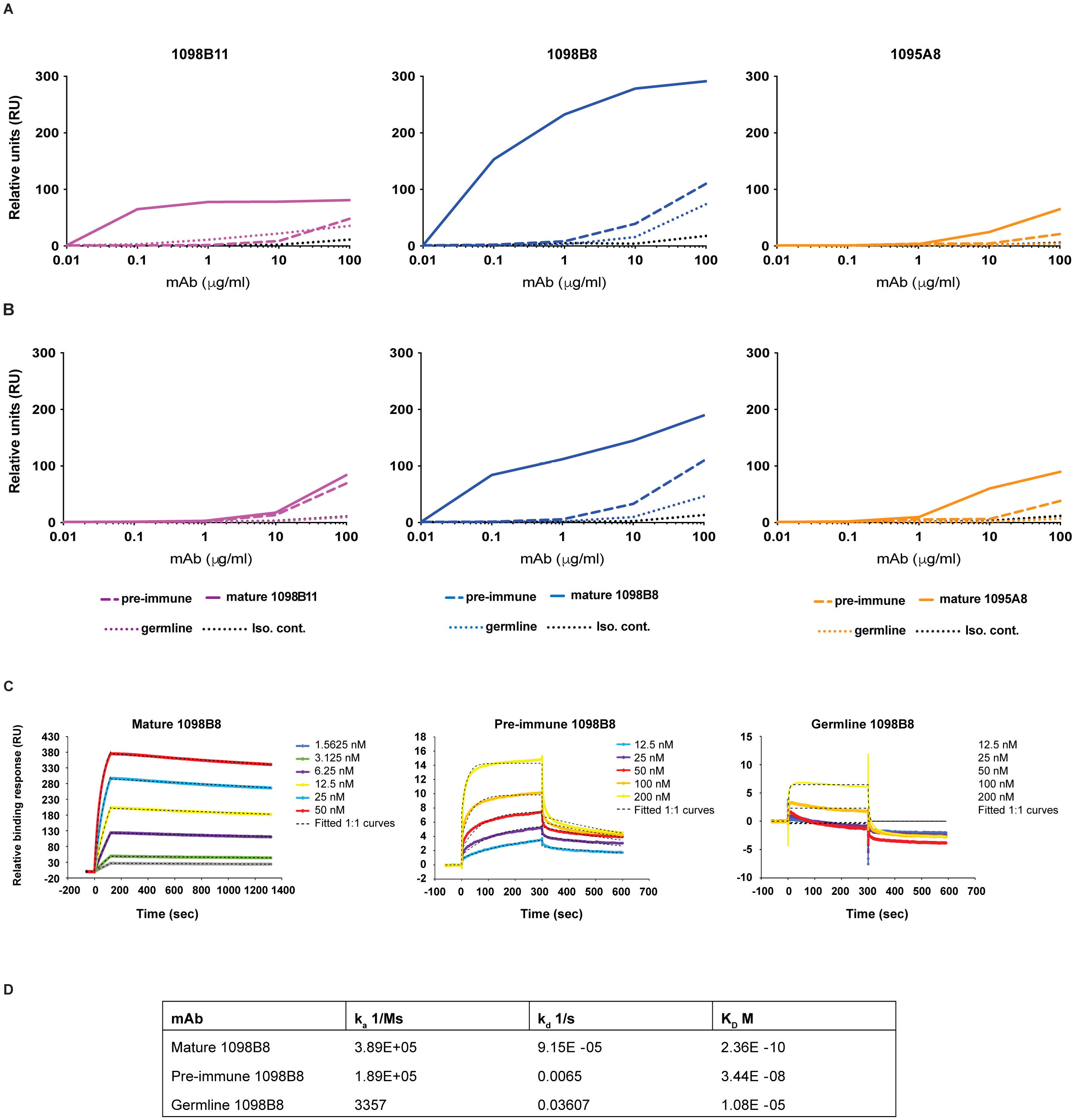
gp120 binding by pre-immune antibodies. (A) Reactivity of germline and pre- and post-vaccination versions of representative mAbs to gp120 determined by ELISA (n = 3 replicates per dilution). Each line represents the mean of an individual mAb. (B) Binding profile of germline and pre- and post-vaccination versions of 1098B8 lineage to gp120 determined by surface plasmon resonance (SPR) analysis and (C) their dissociation constants (K_D_).

## Discussion

While HIV Env is immunogenic and its delivery in numerous forms such as plasmid DNA, protein, and viral vector readily induces Env-specific antibody and B cells; only a minority subset of the responding repertoire is typically able to target the most consequential epitopes or have the potential for conferring protection. As HIV vaccine development progresses, a major emphasis has been placed in trying to focus the humoral vaccine response to highly conserved epitopes and elicit clonal lineages that preferentially target these epitopes. To aid in such nuanced HIV vaccine development efforts we sought to characterize the state and features of gp120-specific B cell repertoire prior to immunization, and the features of those pre-immune B cell clonal lineages that ultimately respond to the vaccine. We determined that in humans a major proportion of the vaccine induced gp120-specific BCR repertoire originates from non-naïve B cells present prior to immunization.

Most of the pre-immune gp120-specific repertoire consisted of IgM (∼60%), however IgA and IgG comprised, ∼25% and ∼15% respectively. Half of the gp120-specific mAb lineages we were able to track back to pre-immune included pre-immune IgA clonal members. While the functional relevance and consequences of the IgA compartment as a major origin of gp120-vaccine induced responses is to be determined, it is consistent with past findings during early HIV infection and exposure that an IgA response to predominantly gp41 but also gp120 (Amos et al., 2015; Ruiz et al., 2016; Yates et al., 2013) is observed particularly in mucosa (Granados-Gonzalez et al., 2008; Seaton et al., 2014). IgA antibodies and B cells are major contributors to mucosal protection, and although we did not look in-depth at the mucosal BCR repertoire, IgA members of the 1131D12 lineage were evident in tonsil after vaccination, and IgA pre-immune members were present in peripheral blood, which may suggest a mucosal association with the origin of this lineage. The systemic and mucosal IgA BCR repertoire is strongly influenced by B1 B cells (Budeus et al., 2021; Kroese et al., 1989), which are an unconventional B cell population that responds quickly against mucosal pathogens through innate-like recognition via IgM, but also that can further develop into highly antigen-specific IgG and IgA B cells (Beagley et al., 1995; Kroese *et al*., 1989; Shikina et al., 2004) indeed, about half of the mucosal IgA has a B1 origin (Budeus *et al*., 2021; Kroese *et al*., 1989). Together this suggests that further assessment of the contribution of mucosal IgA and B1 B cells to the pre-immune gp120 compartment is warranted.

An enrichment of long CDRH3 and increased somatic hypermutation in FRH1 and FRH2 was evident in the pre-immune gp120 compartment, which suggest intrinsic germline and acquired molecular features that determine recognition of gp120. This is consistent with the observation that long CDRH3 are frequently found in responses to HIV infection and vaccination, particularly with Abs targeting gp120 (Easterhoff *et al*., 2017; Wu and Kong, 2016). Additionally, there have been reports of acquired FRH1 and FRH2 motifs that may be conserved amongst public clonotypes recognizing gp120 (Henderson et al., 2019; Karray et al., 1998; Scheid et al., 2011; Zhou et al., 2020).

VH1-2 and VH1-46 usage among the gp120 repertoire is particularly notable, given their use among VRC01-class antibodies which target the CD4 binding site of gp120 and have potent and broad neutralizing activity. Although VH1-2 and VH1-46 were not over-represented in the gp120 pre-immune compartment, they were over-represented among the pre-immune lineages of naïve origin that responded to the immunization. In contrast, VH1-69 which is known to be frequent in antiviral responses (Chen et al., 2019) and VH3-23 which is the most common VH gene used in the human Ig repertoire (Kraj et al., 1997; Wu et al., 2011) was over-represented in those vaccine responding pre-immune lineages of non-naïve origin. It is anticipated that tuning vaccines to elicit VH-specific/class type responses, such as being done with the “germline-targeting” approaches may be more consistent and reproducible amongst vaccinees when targeting the naïve pre-immune compartment, and subsequently more uniform, as being dependent on just germline representations. However, induction of VH-specific/class type responses that are over-represented in the non-naïve pre-immune compartment (e.g. VH1-69) would likely be more inconsistent and highly variable among vaccinees, and heavily dependent on exposure history to the cross-reactive non-HIV antigens that stimulated the initial naïve B cell and the acquisition of cross-reactivity to gp120. Such a process would likely be difficult to reverse engineer into a vaccination strategy. Due to the infrequent nature of gp120-specificity in the pre-immune repertoire and limitations of deep sequencing approaches, highly quantitative tracking and characterization of the lineages that respond or don’t respond to immunization is difficult but should be pursued further given the fundamental heterogeneity we have defined here.

We observed that the non-naïve pre-immune VH (e.g. 1098B8, 1095A8) conferred greater gp120 binding activity than the germline VH. This is consistent with previous demonstration that many HIV bNAbs including 3BNC60, PGT145, CAP256-VRC26, NIH45-46, b12, and 2G12 when reverted to germline have little to no recognition of unmodified gp120 (Andrabi et al., 2015; Gorman et al., 2016; Hoot *et al*., 2013; Scheid *et al*., 2011; Xiao et al., 2009a; Xiao *et al*., 2009b). This supports the possibility that at least a substantial proportion of the non-naïve pre-immune gp120-specific compartment acquired affinity for gp120 through the affinity maturation process against another antigen. Through this process, these clonal lineages and their memory B cells would have likely expanded beyond naïve frequencies, which could have serious implications for immunodominance, and provide rationale for the substantial representation of non-naïve origin lineages in the post-vaccine gp120-specfic repertoire. Several gp120 reactive antibodies have demonstrated cross-reactivity with bacterial antigens (Han et al., 2017; Jeffries et al., 2016; Williams *et al*., 2015), but the sources and dynamics of these cross-reactive antigens remains to be fully resolved. If these initiating antigens or microbes could be identified they may be highly beneficial to use in a pre-priming approach to establish the non-naïve pre-immune gp120 compartment, readied to respond to gp120 immunogens. The current study was not able to resolve these initiating antigens and is a major emphasis of ongoing efforts.

In RV144, the decreased risk of infection observed in vaccinated participants was strongly correlated with the gp120-specific Ab response but protection did not appear to be long-lived, highlighting that durability is a critical element of effective HIV vaccine. We have previously demonstrated that the HVTN 105 vaccine regimen induces long-lived bone marrow plasma cells (Basu *et al*., 2020), the major source of systemic plasma antibody, and here we observe gp120-specific mAb lineage members persisting in the bone marrow for both lineages that arise from naïve (1098B11, 1131A5) and non-naïve (1098B8, 1114G7, 1131D12, 1095A8) origins. Although we cannot exclude some pre-immune seeding of the bone marrow with these non-naïve origin lineage members, this suggests that origin of the gp120 humoral response, whether naïve or non-naïve does not grossly hinder the ability to establish long-lived gp120-specific plasma cells.

In conclusion, we have demonstrated the pool of B cells that respond to initial immunization with HIV Env includes a substantial proportion of antigen-experienced non-naïve B cells. Further defining the mechanisms by which the pre-immune Env-reactive B cell compartment develops and responds to vaccination will likely be necessary for developing broadly efficacious HIV vaccines and may provide insights for other viral vaccines seeking to induce precise Ab classes or germlines.

## Acknowledgements

We are grateful for the assistance of the Rochester Victory Alliance and its staff. We are grateful for the assistance of the University of Rochester Flow Cytometry Core Facility, and the University of Alabama at Birmingham Genomics Core Lab. We are most grateful for the participation of the HVTN 105 study volunteers, and thank the HVTN protocol team for their support of this project. We are very appreciative of the assistance provided by the HIV Vaccine Trials Network in enabling and coordinating this project.

This research was partially funded by the National Institutes of Health [5R21AI116285, 5R01AI117787, 5R01DE027245 to J.J.K., UM1AI069511 to M.C.K.], the University of Rochester Center for AIDS Research P30AI078498 (NIH/NIAID), the University of Alabama at Birmingham Center for AIDS Research P30AI027767 (NIH/NIAID), and the University of Alabama at Birmingham HIV Clinical Trials Unit UM1AI069452 (NIH/NIAID).

## Author Contributions

M.B., A.F.R., J.L., L.X.M, C.A.B., M.C.K., and J.J.K. conceived and designed the study. J.L. and L.X.M. provided critical samples. M.B. and M.S.P. performed the experiments. M.B., M.S.P., C.F.F., A.F.R., and J.J.K. performed the data curation and analysis. M.B. and J.J.K. drafted and edited the manuscript.

## Methods

### Study participants

Blood samples for this study were obtained from 21 participants at the University of Rochester who participated in the HVTN 105 phase 1 randomized, blinded, multisite HIV vaccine clinical trial (ClinicalTrials.gov NCT02207920) (Rouphael *et al*., 2019). All procedures used in this study were approved by the Research Subjects Review Board at the University of Rochester Medical Center and all participants provided written informed consent. All participants were seronegative for HIV infection at the time of enrollment for the study. Participants received different combinations of AIDSVAX^®^ B/E (HIV envelope gp120 of clade B (MN) and E (A244)), DNA-HIV-PT123 (3 plasmids containing DNA of clade C ZM96 gag, clade C ZM96 gp140 and clade C CN54 pol-nef) and placebo administered intra-muscularly at 4 time points over a period of 6 months (Figure 1B). Peripheral blood mononuclear cells (PBMCs) were obtained prior to the vaccination and 7 days post final vaccination. Bone marrow and tonsil biopsy samples collected 7 months post final vaccination.

### HIV-specific antibody secreting cells (ASC) ELISpot

The frequency of HIV-specific ASC in total PBMC were determined by ELISpot similar to as previously described (Basu *et al*., 2020; Kobie et al., 2012; Kobie et al., 2011). Briefly, sterile 96-well PVDF membrane plates (MilliporeSigma, USA) were coated overnight at 4°C with 50 μl of 5 μg/ml HIV-1 Env gp140 SF162.B (NIH AIDS Reagent Program) or 1 μg/ml anti-human IgG or IgM (Jackson Immunoresearch, West Grove, PA) in PBS. Plates were blocked with RPMI1640 (Corning, VA, USA) media with 10 % fetal bovine serum (Atlanta Biologicals, GA, USA) for 2 h at 37°C. Then PBMCs were added in a final volume of 200 μl per well in triplicate. The plates were incubated for ∼ 40 hours at 37 °C in 5% CO_2_ and then washed with PBS containing 0.1% Tween 20. Bound antibodies were detected with 50 μl of 1 μg/ml alkaline phosphatase-conjugated anti-human IgA and horseradish peroxidase-conjugated anti-human IgG (diluted in PBS containing 0.1% Tween 20 and 1% BSA) antibody (Jackson Immunoresearch, West Grove, PA) for 2 hours at 37°C in 5% CO_2_ and then developed with VECTOR Blue, Alkaline Phosphatase Substrate Kit III (Vector Laboratories, Burlingame, CA). The spots per well were counted using the CTL immunospot reader (Cellular Technologies Ltd., Shaker Heights, OH, USA).

### VH Next-Generation Sequencing

For the Ig VH sequencing library preparation, PBMCs were collected prior to the start of the vaccination trial (at M0). gp120+ve and gp120-ve B cell fractions were isolated by magnetic selection using biotinylated HIV Env protein gp120 MN.B and A244.AE. PBMCs were also collected at multiple time points (M1 = 7 days post 2^nd^ vaccination, M3 = 7 days post 3^rd^ vaccination, M6 = 7 days post 4^th^ vaccination) over the course of vaccination. The bone marrow (CD138+ fraction, CD138-fraction or total) and peripheral blood B cells were collected ∼7 months post final vaccination. The Ig heavy chain libraries were prepared as described in Piepenbrink et al., 2021 (Piepenbrink et al., 2021) with modifications. Briefly, total RNA was isolated using the RNeasy Mini Kit (Qiagen, Germany) and treated with DNase I (Turbo DNA-free Kit, Invitrogen, Lithuania). Approximately 1/9 part of this RNA was used for cDNA synthesis in a 20 μL reaction with the qScript cDNA synthesis kit (QuantaBio, MA, USA). Two rounds of PCRs were carried out to generate the libraries with a cocktail of VH1-6 forward primers and IgA, IgG and IgM reverse primers (Basu *et al*., 2020) with the modifications adapted for Illumina Nextera approach. Following PCR, amplicons were analyzed in 1 % agarose gel and bands corresponding to approximately 600 bp were purified with QiAquick gel extraction kit (Qiagen, USA). All the reactions were carried out in triplicates and pooled together during the gel purification. The individual libraries were further purified using the ProNex size-selective purification system (Promega) to select products between 500-700 bp range. Final products were submitted to the University of Alabama at Birmingham Genomics Core Lab, where quality control and quantification were performed using qRT-PCR. Finally, the libraries were pooled together in equimolar ratio and sequenced on an Illumina MiSeq system (Illumina, Inc., CA, USA) using 2 × 300 bp paired-end kits (Illumina MiSeq Reagent Kit v3, 600-cycle, Illumina Inc., CA, USA).

### Sequence analysis of the repertoire

Sequence analysis was performed using an in-house custom analysis pipeline described previously (Basu *et al*., 2020; Nogales et al., 2018; Tipton et al., 2015). Briefly, all sequences were aligned with www.imgt.org/HighV-QUEST (Aouinti et al., 2016) following quality filtering and paired-read joining. To normalize inter-sample variations, 130K sequences were randomly chosen from each library and then analyzed for this study. All clonal lineage assignments were based on identical VH and JH regions, identical HCDR3 length and > 85% HCDR3 nucleotide homology. Expanded clones were determined by plotting size-ranked lineages (% partitions of total sequences) stacked on along the y-axis, and the normalized size per lineage (% of total sequences) plotted on the x-axis. The “knee” of the resulting curve indicates the lineages above which are considered “expanded”. This knee is determined by drawing a line from the top of the curve to the plot origin, and then identifying the longest perpendicular line to the clonality curve.

For lineage tree construction, mAb sequences were parsed against the libraries and the trees were generated with sequences having same V and J gene usage, HCDR3 length, and >80% HCDR3 similariies to the mAb sequences. Within a lineage, instances of only a single identical sequence (for 1098B11 and 1095A8) or only two or less identical sequences (for 1098B8) were removed, with the exception of sequences that match any M0 or M13 or inferred node sequence. The resulting sequences were analyzed using Phylip’s protpars tool (version 3.69s) (Felsenstein, 2005). The output file was then parsed using in-house custom scripts, collapsing any duplicate sequences into an individual node, and was visualized using Cytoscape (Shannon et al., 2003).

### mAb generation

Mature monoclonal antibodies (mAbs) were cloned out from plasmablasts isolated 7 days post final vaccination as previously described in Basu et al., 2020 (Basu *et al*., 2020). Pre-immune representative lineage heavy chains and predicted germline versions of the heavy and light chains were designed with germline V, J and mature CDR3 sequences were synthesized from gBlocks (IDT). These synthesized V-regions were then used for production of recombinant mAbs as full human IgG1 as described previously (Basu *et al*., 2020; Kobie et al., 2015; Tiller et al., 2008). After 8 days of transfection mAbs were purified from culture supernatant using Magna Protein A beads (Promega, WI, USA).

### ELISA

The reactivity profiles of mAbs against HIV Env proteins were detected by ELISA. Briefly, ELISA plates (Nunc MaxiSorp; Thermo Fisher Scientific, NY, USA) were coated overnight at 4 °C with 50 μl of 0.5 μg/ml HIV Env gp120 (MN.B or A244.AE, NIH AIDS Reagent Program) in PBS and blocked with 3% BSA in PBS for 30 min at room temperature. Plasma samples were tested at 1:2500 dilution and mAbs were tested at 10-fold dilutions (100, 10, 1 and 0.1 μg/ml). 50 μl of the samples (diluted in PBS containing 0.05% Tween 20 or PBST) were added per well in triplicate and incubated for 1 h. The reaction was detected using peroxidase-conjugated anti-human IgG (Jackson Immunoreseach, PA, USA), diluted 1:2000 in PBST. Mean OD values of duplicate test samples were divided by control (PBST) and represented as relative units (RU).

### Surface Plasmon Resonance (SPR)

SPR experiments were performed on a Biacore T200 with a CM5 sensor and human IgG capture kit (Cytiva) at 25^0^C using a HEPES buffered saline (HBS) with 0.005%P20 as the running buffer. Anti-human IgG was capture using the protocol recommended by Cytiva over all four flow paths. Between 535 and 580 RU of each antibody was captured using a contact time of 60s and a 10 ul/min flow rate in a HBS running buffer containing 3 mM EDTA and 0.005% P20. MN.B gp120 (AIDS Reagent Repository) was used as the analyte diluted between 1.6 to 200 nM. gp120MN was injected over flow path 1, 2, 3 and 4 with a contact time of 120s, flow rate of 30 ul/min and a long dissociation of 1200s. Flow path 1 was used as reference while antibodies were bound in flow path 2, 3 or 4. 3M MgCl2 was injected over all four flow paths for regeneration using a contact time of 30s and flow rate of 30 ul/min. 1:1 Langmuir model was used to determine kinetics of each antibody. Because of the low level of binding of gp120 (or the saturation at low gp120MN concentrations, steady state kinetics were attempted for both pre-immune 1098B8 and germline 1098B8 by reducing the captured antibody to ∼120 RU and increasing the contact time of analyte to 300s. The pre-immune 1098B8 mAb did not fit well to 1:1 Langmuir kinetics and the lower binding could only be estimated.

### Statistical Analysis

Statistical analysis of VH deep sequencing data was done with Graphpad Prism v7.0 or Matlab, using unpaired Mann-Whitney test or Wilcoxon matched-pairs singed rank test as appropriate.

## References

Abbott, R.K., Lee, J.H., Menis, S., Skog, P., Rossi, M., Ota, T., Kulp, D.W., Bhullar, D., Kalyuzhniy, O., Havenar-Daughton, C., et al. (2018). Precursor Frequency and Affinity Determine B Cell Competitive Fitness in Germinal Centers, Tested with Germline-Targeting HIV Vaccine Immunogens. Immunity 48, 133–146 e136. 10.1016/j.immuni.2017.11.023.

Ahlers, J.D. (2014). All eyes on the next generation of HIV vaccines: strategies for inducing a broadly neutralizing antibody response. Discov Med 17, 187–199.

Amos, J.D., Himes, J.E., Armand, L., Gurley, T.C., Martinez, D.R., Colvin, L., Beck, K., Overman, R.G., Liao, H.X., Moody, M.A., and Permar, S.R. (2015). Rapid Development of gp120-Focused Neutralizing B Cell Responses during Acute Simian Immunodeficiency Virus Infection of African Green Monkeys. J Virol 89, 9485–9498. 10.1128/JVI.01564-15.

Andrabi, R., Voss, J.E., Liang, C.H., Briney, B., McCoy, L.E., Wu, C.Y., Wong, C.H., Poignard, P., and Burton, D.R. (2015). Identification of Common Features in Prototype Broadly Neutralizing Antibodies to HIV Envelope V2 Apex to Facilitate Vaccine Design. Immunity 43, 959–973.10.1016/j.immuni.2015.10.014.

Aouinti, S., Giudicelli, V., Duroux, P., Malouche, D., Kossida, S., and Lefranc, M.P. (2016). IMGT/StatClonotype for Pairwise Evaluation and Visualization of NGS IG and TR IMGT Clonotype (AA) Diversity or Expression from IMGT/HighV-QUEST. Front Immunol 7, 339. 10.3389/fimmu.2016.00339.

Basu, M., Piepenbrink, M.S., Francois, C., Roche, F., Zheng, B., Spencer, D.A., Hessell, A.J., Fucile, C.F., Rosenberg, A.F., Bunce, C.A., et al. (2020). Persistence of HIV-1 Env-Specific Plasmablast Lineages in Plasma Cells after Vaccination in Humans. Cell Rep Med 1. 10.1016/j.xcrm.2020.100015.

Beagley, K.W., Murray, A.M., McGhee, J.R., and Eldridge, J.H. (1995). Peritoneal cavity CD5 (Bla) B cells: cytokine induced IgA secretion and homing to intestinal lamina propria in SCID mice. Immunol Cell Biol 73, 425–432. 10.1038/icb.1995.66.

Bonsignori, M., Liao, H.X., Gao, F., Williams, W.B., Alam, S.M., Montefiori, D.C., and Haynes, B.F. (2017). Antibody-virus co-evolution in HIV infection: paths for HIV vaccine development. Immunol Rev 275, 145–160. 10.1111/imr.12509.

Briney, B., Sok, D., Jardine, J.G., Kulp, D.W., Skog, P., Menis, S., Jacak, R., Kalyuzhniy, O., de Val, N., Sesterhenn, F., et al. (2016). Tailored Immunogens Direct Affinity Maturation toward HIV Neutralizing Antibodies. Cell 166, 1459–1470 e1411. 10.1016/j.cell.2016.08.005.

Budeus, B., Kibler, A., Brauser, M., Homp, E., Bronischewski, K., Ross, J.A., Gorgens, A., Weniger, M.A., Dunst, J., Kreslavsky, T., et al. (2021). Human Cord Blood B Cells Differ from the Adult Counterpart by Conserved Ig Repertoires and Accelerated Response Dynamics. J Immunol. 10.4049/jimmunol.2100113.

Chen, F., Tzarum, N., Wilson, I.A., and Law, M. (2019). VH1-69 antiviral broadly neutralizing antibodies: genetics, structures, and relevance to rational vaccine design. Curr Opin Virol 34, 149–159. 10.1016/j.coviro.2019.02.004.

Chung, A.W., Kumar, M.P., Arnold, K.B., Yu, W.H., Schoen, M.K., Dunphy, L.J., Suscovich, T.J., Frahm, N., Linde, C., Mahan, A.E., et al. (2015). Dissecting Polyclonal Vaccine-Induced Humoral Immunity against HIV Using Systems Serology. Cell 163, 988–998. 10.1016/j.cell.2015.10.027.

ClinicalTrials.gov (2021). A Phase I Trial to Evaluate the Safety and Immunogenicity of eOD-GT8 60mer Vaccine, Adjuvanted. https://clinicaltrials.gov/ct2/show/NCT03547245.

Corey, L., Gilbert, P.B., Juraska, M., Montefiori, D.C., Morris, L., Karuna, S.T., Edupuganti, S., Mgodi, N.M., deCamp, A.C., Rudnicki, E., et al. (2021). Two Randomized Trials of Neutralizing Antibodies to Prevent HIV-1 Acquisition. N Engl J Med 384, 1003–1014. 10.1056/NEJMoa2031738.

Cram, J.A., Fiore-Gartland, A.J., Srinivasan, S., Karuna, S., Pantaleo, G., Tomaras, G.D., Fredricks, D.N., and Kublin, J.G. (2019). Human gut microbiota is associated with HIV-reactive immunoglobulin at baseline and following HIV vaccination. PLoS One 14, e0225622. 10.1371/journal.pone.0225622.

Davis, C.W., Jackson, K.J.L., McCausland, M.M., Darce, J., Chang, C., Linderman, S.L., Chennareddy, C., Gerkin, R., Brown, S.J., Wrammert, J., et al. (2020). Influenza vaccine-induced human bone marrow plasma cells decline within a year after vaccination. Science 370, 237–241. 10.1126/science.aaz8432.

Easterhoff, D., Moody, M.A., Fera, D., Cheng, H., Ackerman, M., Wiehe, K., Saunders, K.O., Pollara, J., Vandergrift, N., Parks, R., et al. (2017). Boosting of HIV envelope CD4 binding site antibodies with long variable heavy third complementarity determining region in the randomized double blind RV305 HIV-1 vaccine trial. PLoS Pathog 13, e1006182. 10.1371/journal.ppat.1006182.

Edupuganti, S., Mgodi, N., Karuna, S.T., Andrew, P., Rudnicki, E., Kochar, N., deCamp, A., De La Grecca, R., Anderson, M., Karg, C., et al. (2021). Feasibility and Successful Enrollment in a Proof-of-Concept HIV Prevention Trial of VRC01, a Broadly Neutralizing HIV-1 Monoclonal Antibody. J Acquir Immune Defic Syndr. 10.1097/QAI.0000000000002639.

Escolano, A., Steichen, J.M., Dosenovic, P., Kulp, D.W., Golijanin, J., Sok, D., Freund, N.T., Gitlin, A.D., Oliveira, T., Araki, T., et al. (2016). Sequential Immunization Elicits Broadly Neutralizing Anti-HIV-1 Antibodies in Ig Knockin Mice. Cell 166, 1445–1458 e1412. 10.1016/j.cell.2016.07.030.

Felsenstein, J. (2005). PHYLIP (Phylogeny Inference Package) version 3.6. Department of Genome Sciences, University of Washington, Seattle, WA.

Gorman, J., Soto, C., Yang, M.M., Davenport, T.M., Guttman, M., Bailer, R.T., Chambers, M., Chuang, G.Y., DeKosky, B.J., Doria-Rose, N.A., et al. (2016). Structures of HIV-1 Env V1V2 with broadly neutralizing antibodies reveal commonalities that enable vaccine design. Nat Struct Mol Biol 23, 81–90. 10.1038/nsmb.3144.

Granados-Gonzalez, V., Claret, J., Berlier, W., Vincent, N., Urcuqui-Inchima, S., Lucht, F., Defontaine, C., Pinter, A., Genin, C., and Riffard, S. (2008). Opposite immune reactivity of serum IgG and secretory IgA to conformational recombinant proteins mimicking V1/V2 domains of three different HIV type 1 subtypes depending on glycosylation. AIDS Res Hum Retroviruses 24, 289–299. 10.1089/aid.2007.0187.

Han, Q., Williams, W.B., Saunders, K.O., Seaton, K.E., Wiehe, K.J., Vandergrift, N., Von Holle, T.A., Trama, A.M., Parks, R.J., Luo, K., et al. (2017). HIV DNA-Adenovirus Multiclade Envelope Vaccine Induces gp41 Antibody Immunodominance in Rhesus Macaques. J Virol 91. 10.1128/JVI.00923-17.

Havenar-Daughton, C., Sarkar, A., Kulp, D.W., Toy, L., Hu, X., Deresa, I., Kalyuzhniy, O., Kaushik, K., Upadhyay, A.A., Menis, S., et al. (2018). The human naive B cell repertoire contains distinct subclasses for a germline-targeting HIV-1 vaccine immunogen. Sci Transl Med 10. 10.1126/scitranslmed.aat0381.

Haynes, B.F., Gilbert, P.B., McElrath, M.J., Zolla-Pazner, S., Tomaras, G.D., Alam, S.M., Evans, D.T., Montefiori, D.C., Karnasuta, C., Sutthent, R., et al. (2012). Immune-correlates analysis of an HIV-1 vaccine efficacy trial. N Engl J Med 366, 1275–1286. 10.1056/NEJMoa1113425.

Henderson, R., Watts, B.E., Ergin, H.N., Anasti, K., Parks, R., Xia, S.M., Trama, A., Liao, H.X., Saunders, K.O., Bonsignori, M., et al. (2019). Selection of immunoglobulin elbow region mutations impacts interdomain conformational flexibility in HIV-1 broadly neutralizing antibodies. Nat Commun 10, 654. 10.1038/s41467-019-08415-7.

Hoot, S., McGuire, A.T., Cohen, K.W., Strong, R.K., Hangartner, L., Klein, F., Diskin, R., Scheid, J.F., Sather, D.N., Burton, D.R., and Stamatatos, L. (2013). Recombinant HIV envelope proteins fail to engage germline versions of anti-CD4bs bNAbs. PLoS Pathog 9, e1003106. 10.1371/journal.ppat.1003106.

Hraber, P., Seaman, M.S., Bailer, R.T., Mascola, J.R., Montefiori, D.C., and Korber, B.T. (2014). Prevalence of broadly neutralizing antibody responses during chronic HIV-1 infection. AIDS 28, 163–169. 10.1097/QAD.0000000000000106.

Huang, D., Abbott, R.K., Havenar-Daughton, C., Skog, P.D., Al-Kolla, R., Groschel, B., Blane, T.R., Menis, S., Tran, J.T., Thinnes, T.C., et al. (2020). B cells expressing authentic naive human VRC01-class BCRs can be recruited to germinal centers and affinity mature in multiple independent mouse models. Proc Natl Acad Sci U S A 117, 22920–22931. 10.1073/pnas.2004489117.

Jardine, J., Julien, J.P., Menis, S., Ota, T., Kalyuzhniy, O., McGuire, A., Sok, D., Huang, P.S., MacPherson, S., Jones, M., et al. (2013). Rational HIV immunogen design to target specific germline B cell receptors. Science 340, 711–716. 10.1126/science.1234150.

Jardine, J.G., Ota, T., Sok, D., Pauthner, M., Kulp, D.W., Kalyuzhniy, O., Skog, P.D., Thinnes, T.C., Bhullar, D., Briney, B., et al. (2015). HIV-1 VACCINES. Priming a broadly neutralizing antibody response to HIV-1 using a germline-targeting immunogen. Science 349, 156–161. 10.1126/science.aac5894.

Jeffries, T.L., Jr., Sacha, C.R., Pollara, J., Himes, J., Jaeger, F.H., Dennison, S.M., McGuire, E., Kunz, E., Eudailey, J.A., Trama, A.M., et al. (2016). The function and affinity maturation of HIV-1 gp120-specific monoclonal antibodies derived from colostral B cells. Mucosal Immunol 9, 414–427. 10.1038/mi.2015.70.

Karasavvas, N., Billings, E., Rao, M., Williams, C., Zolla-Pazner, S., Bailer, R.T., Koup, R.A., Madnote, S., Arworn, D., Shen, X., et al. (2012). The Thai Phase III HIV Type 1 Vaccine trial (RV144) regimen induces antibodies that target conserved regions within the V2 loop of gp120. AIDS Res Hum Retroviruses 28, 1444–1457. 10.1089/aid.2012.0103.

Karray, S., Juompan, L., Maroun, R.C., Isenberg, D., Silverman, G.J., and Zouali, M. (1998). Structural basis of the gp120 superantigen-binding site on human immunoglobulins. J Immunol 161, 6681–6688.

Kobie, J.J., Alcena, D.C., Zheng, B., Bryk, P., Mattiacio, J.L., Brewer, M., Labranche, C., Young, F.M., Dewhurst, S., Montefiori, D.C., et al. (2012). 9G4 autoreactivity is increased in HIV-infected patients and correlates with HIV broadly neutralizing serum activity. PLoS One 7, e35356. 10.1371/journal.pone.0035356.

Kobie, J.J., Zheng, B., Bryk, P., Barnes, M., Ritchlin, C.T., Tabechian, D.A., Anandarajah, A.P., Looney, R.J., Thiele, R.G., Anolik, J.H., et al. (2011). Decreased influenza-specific B cell responses in rheumatoid arthritis patients treated with anti-tumor necrosis factor. Arthritis Res Ther 13, R209. 10.1186/ar3542.

Kobie, J.J., Zheng, B., Piepenbrink, M.S., Hessell, A.J., Haigwood, N.L., Keefer, M.C., and Sanz, I. (2015). Functional and Molecular Characteristics of Novel and Conserved Cross-Clade HIV Envelope Specific Human Monoclonal Antibodies. Monoclon Antib Immunodiagn Immunother 34, 65–72. 10.1089/mab.2014.0064.

Kraj, P., Rao, S.P., Glas, A.M., Hardy, R.R., Milner, E.C., and Silberstein, L.E. (1997). The human heavy chain Ig V region gene repertoire is biased at all stages of B cell ontogeny, including early pre-B cells. J Immunol 158, 5824–5832.

Krebs, S.J., Kwon, Y.D., Schramm, C.A., Law, W.H., Donofrio, G., Zhou, K.H., Gift, S., Dussupt, V., Georgiev, I.S., Schatzle, S., et al. (2019). Longitudinal Analysis Reveals Early Development of Three MPER-Directed Neutralizing Antibody Lineages from an HIV-1-Infected Individual. Immunity 50, 677–691 e613. 10.1016/j.immuni.2019.02.008.

Kroese, F.G., Butcher, E.C., Stall, A.M., Lalor, P.A., Adams, S., and Herzenberg, L.A. (1989). Many of the IgA producing plasma cells in murine gut are derived from self-replenishing precursors in the peritoneal cavity. Int Immunol 1, 75–84. 10.1093/intimm/1.1.75.

Lin, Y.R., Parks, K.R., Weidle, C., Naidu, A.S., Khechaduri, A., Riker, A.O., Takushi, B., Chun, J.H., Borst, A.J., Veesler, D., et al. (2020). HIV-1 VRC01 Germline-Targeting Immunogens Select Distinct Epitope-Specific B Cell Receptors. Immunity 53, 840–851 e846. 10.1016/j.immuni.2020.09.007.

Mgodi, N.M., Takuva, S., Edupuganti, S., Karuna, S., Andrew, P., Lazarus, E., Garnett, P., Shava, E., Mukwekwerere, P.G., Kochar, N., et al. (2021). A phase 2b study to evaluate the safety and efficacy of VRC01 broadly neutralizing monoclonal antibody in reducing acquisition of HIV-1 infection in women in sub-Saharan Africa: baseline findings. J Acquir Immune Defic Syndr. 10.1097/QAI.0000000000002649.

Moody, M.A., Yates, N.L., Amos, J.D., Drinker, M.S., Eudailey, J.A., Gurley, T.C., Marshall, D.J., Whitesides, J.F., Chen, X., Foulger, A., et al. (2012). HIV-1 gp120 vaccine induces affinity maturation in both new and persistent antibody clonal lineages. J Virol 86, 7496–7507. 10.1128/JVI.00426-12.

Mouquet, H., and Nussenzweig, M.C. (2012). Polyreactive antibodies in adaptive immune responses to viruses. Cell Mol Life Sci 69, 1435–1445. 10.1007/s00018-011-0872-6.

Ng’uni, T., Chasara, C., and Ndhlovu, Z.M. (2020). Major Scientific Hurdles in HIV Vaccine Development: Historical Perspective and Future Directions. Front Immunol 11, 590780. 10.3389/fimmu.2020.590780.

Nogales, A., Piepenbrink, M.S., Wang, J., Ortega, S., Basu, M., Fucile, C.F., Treanor, J.J., Rosenberg, A.F., Zand, M.S., Keefer, M.C., et al. (2018). A Highly Potent and Broadly Neutralizing H1 Influenza-Specific Human Monoclonal Antibody. Sci Rep 8, 4374. 10.1038/s41598-018-22307-8.

Piepenbrink, M.S., Park, J.G., Oladunni, F.S., Deshpande, A., Basu, M., Sarkar, S., Loos, A., Woo, J., Lovalenti, P., Sloan, D., et al. (2021). Therapeutic activity of an inhaled potent SARS-CoV-2 neutralizing human monoclonal antibody in hamsters. Cell Rep Med 2, 100218. 10.1016/j.xcrm.2021.100218.

Planchais, C., Kok, A., Kanyavuz, A., Lorin, V., Bruel, T., Guivel-Benhassine, F., Rollenske, T., Prigent, J., Hieu, T., Prazuck, T., et al. (2019). HIV-1 Envelope Recognition by Polyreactive and Cross-Reactive Intestinal B Cells. Cell Rep 27, 572–585 e577. 10.1016/j.celrep.2019.03.032.

Rerks-Ngarm, S., Pitisuttithum, P., Nitayaphan, S., Kaewkungwal, J., Chiu, J., Paris, R., Premsri, N., Namwat, C., de Souza, M., Adams, E., et al. (2009). Vaccination with ALVAC and AIDSVAX to prevent HIV-1 infection in Thailand. N Engl J Med 361, 2209–2220. 10.1056/NEJMoa0908492.

Rouphael, N.G., Morgan, C., Li, S.S., Jensen, R., Sanchez, B., Karuna, S., Swann, E., Sobieszczyk, M.E., Frank, I., Wilson, G.J., et al. (2019). DNA priming and gp120 boosting induces HIV-specific antibodies in a randomized clinical trial. J Clin Invest 129, 4769–4785. 10.1172/JCI128699.

Ruiz, M.J., Ghiglione, Y., Falivene, J., Laufer, N., Holgado, M.P., Socias, M.E., Cahn, P., Sued, O., Giavedoni, L., Salomon, H., et al. (2016). Env-Specific IgA from Viremic HIV-Infected Subjects Compromises Antibody-Dependent Cellular Cytotoxicity. J Virol 90, 670–681. 10.1128/JVI.02363-15.

Scheid, J.F., Mouquet, H., Ueberheide, B., Diskin, R., Klein, F., Oliveira, T.Y., Pietzsch, J., Fenyo, D., Abadir, A., Velinzon, K., et al. (2011). Sequence and structural convergence of broad and potent HIV antibodies that mimic CD4 binding. Science 333, 1633–1637. 10.1126/science.1207227.

Seaton, K.E., Ballweber, L., Lan, A., Donathan, M., Hughes, S., Vojtech, L., Moody, M.A., Liao, H.X., Haynes, B.F., Galloway, C.G., et al. (2014). HIV-1 specific IgA detected in vaginal secretions of HIV uninfected women participating in a microbicide trial in Southern Africa are primarily directed toward gp120 and gp140 specificities. PLoS One 9, e101863. 10.1371/journal.pone.0101863.

Setliff, I., McDonnell, W.J., Raju, N., Bombardi, R.G., Murji, A.A., Scheepers, C., Ziki, R., Mynhardt, C., Shepherd, B.E., Mamchak, A.A., et al. (2018). Multi-Donor Longitudinal Antibody Repertoire Sequencing Reveals the Existence of Public Antibody Clonotypes in HIV-1 Infection. Cell Host Microbe 23, 845–854 e846. 10.1016/j.chom.2018.05.001.

Shannon, P., Markiel, A., Ozier, O., Baliga, N.S., Wang, J.T., Ramage, D., Amin, N., Schwikowski, B., and Ideker, T. (2003). Cytoscape: a software environment for integrated models of biomolecular interaction networks. Genome Res 13, 2498–2504. 10.1101/gr.1239303.

Shikina, T., Hiroi, T., Iwatani, K., Jang, M.H., Fukuyama, S., Tamura, M., Kubo, T., Ishikawa, H., and Kiyono, H. (2004). IgA class switch occurs in the organized nasopharynx-and gut-associated lymphoid tissue, but not in the diffuse lamina propria of airways and gut. J Immunol 172, 6259–6264. 10.4049/jimmunol.172.10.6259.

Soto, C., Bombardi, R.G., Branchizio, A., Kose, N., Matta, P., Sevy, A.M., Sinkovits, R.S., Gilchuk, P., Finn, J.A., and Crowe, J.E., Jr. (2019). High frequency of shared clonotypes in human B cell receptor repertoires. Nature 566, 398–402. 10.1038/s41586-019-0934-8.

Tiller, T., Meffre, E., Yurasov, S., Tsuiji, M., Nussenzweig, M.C., and Wardemann, H. (2008). Efficient generation of monoclonal antibodies from single human B cells by single cell RT-PCR and expression vector cloning. J Immunol Methods 329, 112–124. 10.1016/j.jim.2007.09.017.

Tipton, C.M., Fucile, C.F., Darce, J., Chida, A., Ichikawa, T., Gregoretti, I., Schieferl, S., Hom, J., Jenks, S., Feldman, R.J., et al. (2015). Diversity, cellular origin and autoreactivity of antibody-secreting cell population expansions in acute systemic lupus erythematosus. Nat Immunol 16, 755–765. 10.1038/ni.3175.

Trama, A.M., Moody, M.A., Alam, S.M., Jaeger, F.H., Lockwood, B., Parks, R., Lloyd, K.E., Stolarchuk, C., Scearce, R., Foulger, A., et al. (2014). HIV-1 envelope gp41 antibodies can originate from terminal ileum B cells that share cross-reactivity with commensal bacteria. Cell Host Microbe 16, 215–226. 10.1016/j.chom.2014.07.003.

West, A.P., Jr., Diskin, R., Nussenzweig, M.C., and Bjorkman, P.J. (2012). Structural basis for germ-line gene usage of a potent class of antibodies targeting the CD4-binding site of HIV-1 gp120. Proc Natl Acad Sci U S A 109, E2083–2090. 10.1073/pnas.1208984109.

WHO (2020). WHO HIV/AIDS Data and Statistics. https://www.who.int/hiv/data/en.

Wiehe, K., Easterhoff, D., Luo, K., Nicely, N.I., Bradley, T., Jaeger, F.H., Dennison, S.M., Zhang, R., Lloyd, K.E., Stolarchuk, C., et al. (2014). Antibody light-chain-restricted recognition of the site of immune pressure in the RV144 HIV-1 vaccine trial is phylogenetically conserved. Immunity 41, 909–918. 10.1016/j.immuni.2014.11.014.

Williams, W.B., Han, Q., and Haynes, B.F. (2018). Cross-reactivity of HIV vaccine responses and the microbiome. Curr Opin HIV AIDS 13, 9–14. 10.1097/COH.0000000000000423.

Williams, W.B., Liao, H.X., Moody, M.A., Kepler, T.B., Alam, S.M., Gao, F., Wiehe, K., Trama, A.M., Jones, K., Zhang, R., et al. (2015). HIV-1 VACCINES. Diversion of HIV-1 vaccine-induced immunity by gp41-microbiota cross-reactive antibodies. Science 349, aab1253. 10.1126/science.aab1253.

Wu, X., and Kong, X.P. (2016). Antigenic landscape of the HIV-1 envelope and new immunological concepts defined by HIV-1 broadly neutralizing antibodies. Curr Opin Immunol 42, 56–64. 10.1016/j.coi.2016.05.013.

Wu, X., Zhang, Z., Schramm, C.A., Joyce, M.G., Kwon, Y.D., Zhou, T., Sheng, Z., Zhang, B., O’Dell, S., McKee, K., et al. (2015). Maturation and Diversity of the VRC01-Antibody Lineage over 15 Years of Chronic HIV-1 Infection. Cell 161, 470–485. 10.1016/j.cell.2015.03.004.

Wu, Y.C., Kipling, D., and Dunn-Walters, D.K. (2011). The relationship between CD27 negative and positive B cell populations in human peripheral blood. Front Immunol 2, 81. 10.3389/fimmu.2011.00081.

Xiao, X., Chen, W., Feng, Y., and Dimitrov, D.S. 2009a). Maturation Pathways of Cross-Reactive HIV-1 Neutralizing Antibodies. Viruses 1, 802–817. 10.3390/v1030802.

Xiao, X., Chen, W., Feng, Y., Zhu, Z., Prabakaran, P., Wang, Y., Zhang, M.Y., Longo, N.S., and Dimitrov, D.S. (2009b). Germline-like predecessors of broadly neutralizing antibodies lack measurable binding to HIV-1 envelope glycoproteins: implications for evasion of immune responses and design of vaccine immunogens. Biochem Biophys Res Commun 390, 404–409. 10.1016/j.bbrc.2009.09.029.

Yates, N.L., Liao, H.X., Fong, Y., deCamp, A., Vandergrift, N.A., Williams, W.T., Alam, S.M., Ferrari, G., Yang, Z.Y., Seaton, K.E., et al. (2014). Vaccine-induced Env V1-V2 IgG3 correlates with lower HIV-1 infection risk and declines soon after vaccination. Sci Transl Med 6, 228ra239. 10.1126/scitranslmed.3007730.

Yates, N.L., Stacey, A.R., Nolen, T.L., Vandergrift, N.A., Moody, M.A., Montefiori, D.C., Weinhold, K.J., Blattner, W.A., Borrow, P., Shattock, R., et al. (2013). HIV-1 gp41 envelope IgA is frequently elicited after transmission but has an initial short response half-life. Mucosal Immunol 6, 692–703. 10.1038/mi.2012.107.

Zhou, J.O., Zaidi, H.A., Ton, T., and Fera, D. (2020). The Effects of Framework Mutations at the Variable Domain Interface on Antibody Affinity Maturation in an HIV-1 Broadly Neutralizing Antibody Lineage. Front Immunol 11, 1529. 10.3389/fimmu.2020.01529.

Zolla-Pazner, S., deCamp, A., Gilbert, P.B., Williams, C., Yates, N.L., Williams, W.T., Howington, R., Fong, Y., Morris, D.E., Soderberg, K.A., et al. (2014a). Vaccine-induced IgG antibodies to V1V2 regions of multiple HIV-1 subtypes correlate with decreased risk of HIV-1 infection. PLoS One 9, e87572. 10.1371/journal.pone.0087572.

Zolla-Pazner, S., deCamp, A.C., Cardozo, T., Karasavvas, N., Gottardo, R., Williams, C., Morris, D.E., Tomaras, G., Rao, M., Billings, E., et al. (2013). Analysis of V2 antibody responses induced in vaccinees in the ALVAC/AIDSVAX HIV-1 vaccine efficacy trial. PLoS One 8, e53629. 10.1371/journal.pone.0053629.

Zolla-Pazner, S., Edlefsen, P.T., Rolland, M., Kong, X.P., deCamp, A., Gottardo, R., Williams, C., Tovanabutra, S., Sharpe-Cohen, S., Mullins, J.I., et al. (2014b). Vaccine-induced Human Antibodies Specific for the Third Variable Region of HIV-1 gp120 Impose Immune Pressure on Infecting Viruses. EBioMedicine 1, 37–45. 10.1016/j.ebiom.2014.10.022.

